# Co-Stimulatory Blockade Prevents Intragraft Accrual of Class-Switched, Activated B Cells Despite Failing to Prevent T-Cell Mediated Rejection

**DOI:** 10.1101/2025.10.24.683156

**Authors:** J. Timothy Caldwell, Tiffany Shi, Ashley R. Burg, Ashley Koby, Krishna M. Roskin, Anna L. Peters, Erica A.K. DePasquale, E. Steve Woodle, David A. Hildeman

**Author notes:** Corresponding authors: David A Hildeman, Cincinnati Children’s Hospital Medical Center, 3333 Burnet Ave, Cincinnati, OH 45229, USA., E. Steve Woodle, University of Cincinnati College of Medicine, 3230 Eden Ave, Cincinnati, OH 45267, USA. these authors contributed equally and share first authorship (J. Timothy Caldwell, Tiffany Shi). authors contributed equally and share senior authorship (E. Steve Woodle, David A. Hildeman).

## Abstract

Previously, we defined the transcriptomes and clonality of intragraft CD8^+^ T cell during renal allograft rejection. Here, using single cell RNA sequencing (scRNAseq), we investigated non-CD8^+^ immune cells during T-cell-mediated rejection (TCMR) under different maintenance immunosuppression (mIS) regimens: tacrolimus, and co-stimulatory blockade (CoB) with belatacept (CTLA4-Ig) or iscalimab (anti−CD40). Myeloid cells comprised of DCs, monocytes and macrophages whose proportion and gene expression were similar between mIS regimens. Given their transcriptiomic similarities, we analyzed publicly-available scRNAseq/CITEseq datasets as well as immunofluorescence staining to resolve independent subpopulations of γ/δ T cells and NK cells. Intragraft B cells consisted of clusters of naïve, plasmablast, and class-switched B (B_CS_) cells, with the latter being diminished in CoB mIS. Intragraft CD4^+^ T cells consisted of FoxP3^+^ regulatory (Treg), exhausted, Th17, and CXCL13^+^ peripheral helper (Tph) cells whose proportions differed based in mIS, and the latter two had increased clonal expansion. Notably, cell-cell communication analysis indicated Th17 and Tph cell interactions with B_CS_ cells in tacrolimus, but not CoB, samples. Thus, although failing to prevent TCMR, CoB mIS modulates the accrual of CD4^+^ T cells and inhibits the intragraft accrual of B_CS_ cells, possibly reflective of clinical observations of less chronic antibody-mediated rejection under CoB mIS.

## 1. Introduction

Despite advances in immunosuppression (IS), allograft rejection remains a major contributor to kidney allograft loss and subsequently limits patient quality of life and survival^1^. T-cell-mediated rejection (TCMR), which is largely effected by attack from CD8^+^ T cells^2^, independently shortens graft life and also predisposes to subsequent rejections^1,3^. While the introduction of calcineurin inhibitors (CNIs) has greatly reduced the incidence of TCMR, their long-term use as maintenance immunosuppression (mIS) is associated with severe metabolic, renal and cardiovascular toxicities^4^. To avoid such toxicities, targeted mIS approaches with costimulation blockade (CoB) have been developed, including belatacept^5–7^ (CTLA4-Ig) and more recently, anti-CD40 monoclonal antibodies^8,9, 10^. Given the critical role of co-stimulation in both B and T cell function, CoB is an attractive target for mIS as it has fewer long-term multisystem adverse toxicities compared with CNIs^11,12^.

Consistent with its mechanism of action, CoB through either the CD28/B7 or CD40/CD154 pathway has been shown to disrupt germinal centers and prevent both B cell maturation and donor specific antibody (DSA) formation^13–16^. Additionally, T cells require costimulation for successful expansion and maturation after antigen stimulation to prevent anergy^17,18^. CTLA4-Ig blocks interactions between CD28 on T cells and B7 on antigen presenting cells (APC)^5^. While CTLA4-Ig prevents the expansion of naïve T cells^5^, it is less effective at preventing expansion of memory T cells^19^. Similarly, CD154 (also known as CD40L, encoded by *CD40LG* gene, however given recognition of additional CD40 binding partners^20^ will refer to protein as CD154) expressed by T cells is critical for licensing CD40^+^ DCs to promote further T cell activation as well as the expansion and class-switching of CD40^+^ B cells^21–24^. Importantly, one of the potential problems of CoB-mediated mIS is the development of rejection mediated by pre-existing memory T cells, which are not reliant on costimulation and are resistant to CoB ^17,25–27^.

The roles and impacts of most immune cell types at the time of TCMR is only partially understood. The myeloid compartment, for example, has yielded mixed results in terms of being either pro- or anti-rejection, likely due to both experimental model, contrasting temporal functions, and the remarkable cellular complexity and plasticity within innate immune cells^28–30^. Traditionally thought of as important for antibody-mediated rejection (AMR), B cells have been suggested to play a role in TCMR, including as APCs facilitating activation of CD4^+^ T cells^31^.

This has historically been thought to occur in secondary lymphoid organs^32^, however, the development of tertiary lymphoid organs (TLO) in the allograft is also known to occur^33–35^. While these studies have largely focused on chronic rejection, the presence of TLOs suggest a role for intra-graft B cell activation^36^. On the other hand, regulatory B cells have been described to promote a tolerogenic state^37–39^. Further adding to this complexity is that, during chronic rejection, nearly all intragraft B cells appear to be auto-but not allo-reactive^40^. However, little has been done to characterize intragraft B cells during TCMR. The roles of CD4^+^ T cells and γ/δ T cells are similarly complex, with a striking range of reported functions and impacts on the allograft^41,42^. For example, circulating T follicular helper (Tfh) cells have been shown to predict DSA formation and contribute to alloreactivity post-transplant^43,44^, while a unique CD57^+^ CD4 population has been associated with belatacept-resistant rejection^45^. Similarly, we and others have described potential γ/δ T cells within allografts, but their clear delineation and function has remained elusive^46,47^. Nonetheless, an intragraft analysis of non-CD8 immune cells, their potential cross-talk, and the influence of mIS on these cells during human allograft rejection at whole genome scale, remains understudied.

We previously employed single-cell RNA sequencing (scRNAseq) and single-cell TCR sequencing (scTCRseq) to analyze graft-infiltrating CD8^+^ T cell clonality and phenotype during TCMR, demonstrating IS-specific impacts on the transcriptomic landscape of CD8^+^ T cell subpopulations^47^. Here, we analyze the transcriptomic landscape of intragraft non-CD8^+^ immune cells present at the time of TCMR diagnosis under varying mIS regimens. We identify the differences between CD4^+^ T cells and B cells, but not other innate immune cells in patients treated with tacrolimus, belatacept, and iscalimab (anti-CD40^8^) mIS. Taken together, our data show that the type of mIS differentially impacts the B cell and CD4^+^ T cell compartments with implications on mechanisms of rejection, as well as potential subsequent rejection risk.

## 2.0 Methods

### 2.1 Data acquisition

Kidney allograft biopsy single cell sequencing (scRNAseq, scTCRseq) data were derived from 10 participants as reported in our previous study^47^. Briefly, biopsy cores taken at the time of TCMR diagnosis were subjected to cryopreservation, cold-protease digestion, and processing using 10x Chromium 5′ assay with TCR sequencing using the Chromium Next GEM Single Cell V(D)J Reagent Kit v1.1. Genomics data are available on NCBI BioProject, accession number PRJNA974568. Sequencing data from liver rejection samples are accessible from the Gene Expression Omnibus (GEO) using accession GSE256141. Reference datasets from other groups were obtained as follows: NK cells from Jaeger et al. were downloaded from https://github.com/aulezko/HumanILC1^48^; multimodal scRNAseq/CITEseq data from Hao et al. were downloaded from GEO (accession GSE164378) and https://atlas.fredhutch.org/nygc/multimodal-pbmc/^49^; integrated NK data from Rebuffet et al. were downloaded or explored using the interactive explorer at https://collections.cellatlas.io/meta-nk^50^. Visualizations were created as below or the website data explorers.

### 2.2 Single-cell RNASeq analysis pipeline

Primary single-cell analyses of cells from kidney biopsies were carried out with R (version 4.2.0) running inside RStudio (version 4.1.1) using Seurat (version 4.1.0) as previously described^47^. Briefly, samples were integrated following the standard vignette and differential gene expression analyses were performed using Seurat (version 4.1.0)^49^ functions FindAllMarkers or FindMarkers with default settings to identify significantly expressed genes at adjusted p-value of <0.05. Visualization was accomplished using either default Seurat functions or variations from the scCustomize package^51^. Integration with liver biopsy datasets was done using the Harmony package in R^52^ and Seurat v5 (version 5.3.0^53^). Briefly, datasets were merged, scaled, and normalized. The RunHarmony command was used with parameters for “orig.ident” and “dataset” (liver or kidney) to generate harmony embeddings that were then used for UMAP and neighbor calculations.

CellChat analysis was performed using the CellChat Package^54,55^. Clusters were filtered to only include clusters that were present in each sample prior to running analysis. Analysis followed the standard vignette using default settings. After the CellChat object was created and significant interactions were determined, the counts for each cell-cell interaction were extracted. Interactions involving *HLA* genes were excluded from the total, as the interaction of each class II HLA gene with CD4 artificially inflated the number of interactions.

NicheNet analysis was performed using the NicheNet package^56^. A subsetted Seurat object that only contained the cell clusters present in each mIS group was created, and proliferating cells were then excluded. The nichenet_seuratobj_cluster_de function was used using default settings per the standard vignette to determine candidate ligands responsible for differences between groups as indicated in the text. The top 5 ligands from each group as ranked by AUPR were extracted and presented for further analysis.

### 2.3 Immunofluorescence

FFPE kidney tissue slides were subjected to deparaffinization, rehydration, and antigen retrieval with Trilogy® Pretreatment solution (cat. no. 920P-09, Sigma-Aldrich). Two Coplin jars of solution were preheated in boiling water. Slides were placed in one jar for 30 minutes while boiling. They were then transferred to the 2^nd^ hot jar, taken out of the boiling water, and placed on a rocker for 30 minutes. Slides were washed with PBS with rocking and blocked with donkey serum for 1 hour. Next, the primary antibodies incubated overnight at 4°C with rabbit anti-human CD3ε (dilution 1:50, 85061, Cell Signaling Technology), goat anti-human CD56 (dilution 1:50, AF2408, R&D Systems), mouse anti-human TCR δ (dilution 1:50, sc-100289, Santa Cruz), and rat anti-human CD16 (dilution 1:100, ab308607, Abcam), all were diluted in the blocking solution used previously. After washing, secondary antibodies were applied: donkey anti-rabbit Alexa Fluor 488 (dilution 1:50, PIA32790TR, Fisher Scientific); donkey anti-goat Dylight 550 (dilution 1:50, PISA510087, Fisher Scientific); donkey anti-mouse Alexa Fluor 647 (dilution 1:50, PIA32787TR, Fisher Scientific); and donkey anti-rat Dylight 755 (dilution 1:100, PISA510031, Fisher Scientific). All were diluted in PBS and incubated at room temperature for 1 hour. The slides were washed, and TrueBlack® Lipofuscin Autofluorescence Quencher (23007, Biotium) was added for 3 minutes to reduce autofluorescence. Following another washing step, DAPI (62248, Thermo Scientific) was added for 5 minutes. The slides were mounted (P36935, Invitrogen) and cover-slipped. Images were taken on a Nikon AXR inverted confocal laser scanning microscope in the Bio-Imaging and Analysis Facility at Cincinnati Children’s Hospital Medical Center (RRID: SCR_022628).

## 3. Results

### 3.1 Maintenance Immunosuppression has Minimal Effects on Myeloid Cells

Previously, we employed scRNAseq combined with scTCRseq to determine graft-infiltrating CD8^+^ T cell clonality and gene expression profiles during renal allograft rejection under 3 IS modalities: tacrolimus (N=4), belatacept (N=3), and iscalimab (N=3)^47^. We focused on samples undergoing TCMR, as non-rejecting samples had a paucity of immune cells. Here, we analyzed the non-CD8^+^ T cell immune compartment in the index biopsy samples from those 10 patients. We first compuationally segregated the myeloid cell populations and annotated individual cell types. Cluster analysis revealed 9 cell subsets based on their differentially expressed genes (DEG) (Supplementary Table 1), including: plasmacytoid dendritic cells (pDCs), classical monocytes, non-classical monocytes, intermediate monocytes, conventional DCs, monocyte-derived DCs, anti-inflammatory DCs, macrophages, and a very small cluster of inflammatory macrophages (Fig. 1 A, B).

**Figure 1.**
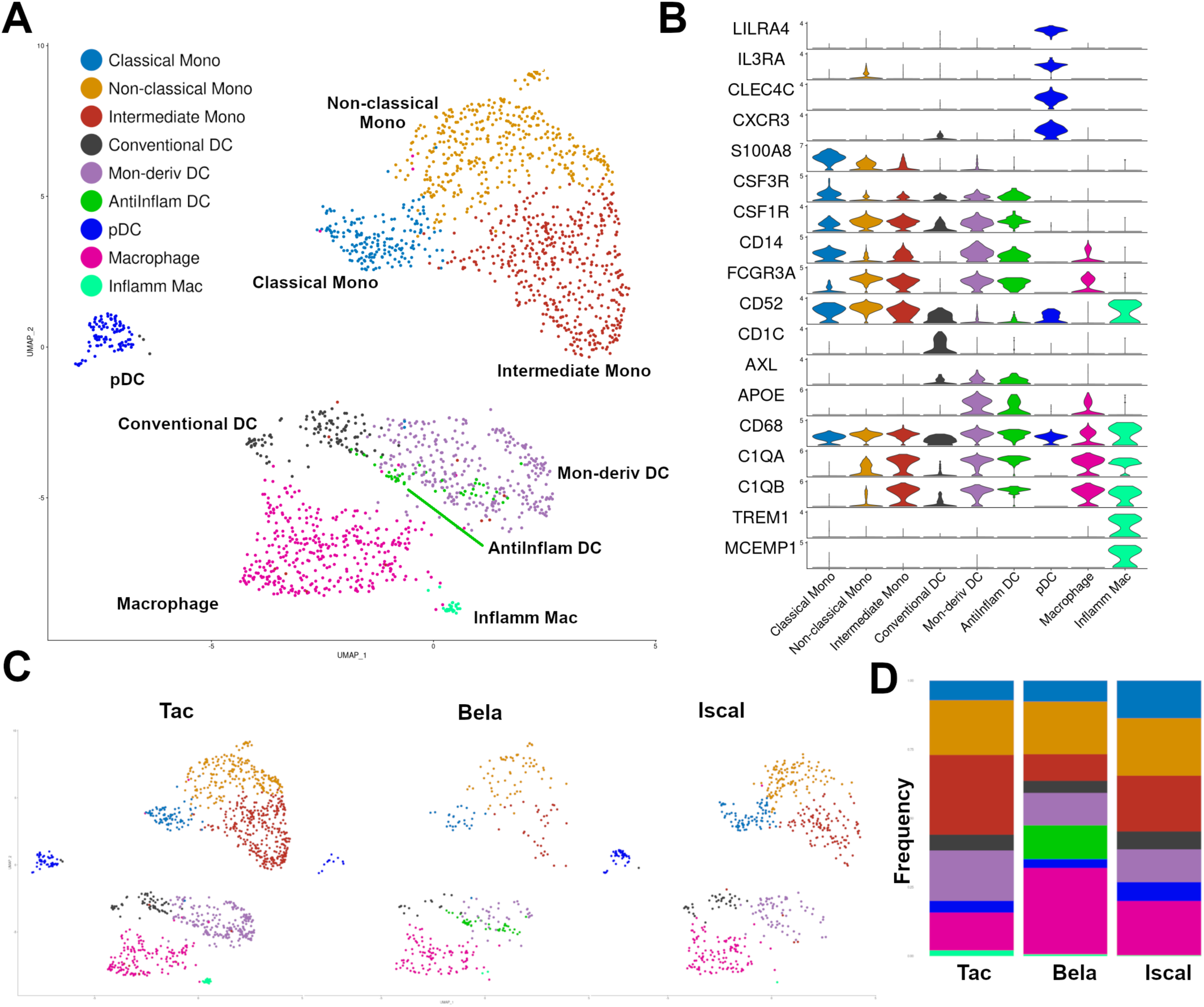
Myeloid populations are preserved across maintenance immunosuppressive regimens. A) UMAP showing distribution and annotation of identified myeloid populations. B) Violin plot showing expression of relevant marker genes by myeloid cluster. C) UMAP showing distribution of cells separated by mIS regimen and presence of an anti-inflammatory DC population. D) Bar graph showing proportional makeup of myeloid population by mIS regimen. DC = dendritic cell; pDC = plasmacytoid dendritic cell; UMAP = uniform manifold approximation and projection; mIS = maintenance immunosuppression.

Most myeloid cell subsets were represented at similar frequencies in biopsy samples from all three IS modalities. (Figures 1C, 1D, S1A). Gene expression within the myeloid compartment revealed surprisingly minimal differences between mIS groups, with most DEG being related to ribosomal or genes associated with protein transcription (Supplementary Table 2). In two belatacept treated patients, we identified a potentially novel cluster of anti-inflammatory DCs (Fig. 1C, D, S1A), and pathway analysis of DEG demonstrated enrichment of immune regulatory pathways, with top two enriched GO:Biological Process being Negative Regulation of Leukocyte Activation (GO:0002695) and Regulation of Chemokine Production (GO:0032642). However, as this population was only uncovered in 2 patients, it is unclear if these cells are more common in patients with belatcept-refractory rejection (BRR) relative to other mIS regimens. Beyond this unique cluster, mIS had little effect on the subsets or gene expression of intragraft myeloid cells in patients undergoing rejection.

### 3.2 ã/ä T Cells Are Present at Time of Rejection and Demonstrate a Range of Transcriptomic Phenotypes

An intriguing finding from our previous work^47^ was the presence of what we deemed γ/δ T cells (based on their clustering near α/β T cells, their expression of *TRDC*, and lack of *CD4/CD8/TRAC/TRBC* expression) in index biopsies of all patients (Figure 2A, S1B). Given their transcriptomic similarity to NK cells and recent work directly comparing the transcriptomes of NK cells and γδ T cells^48,50,57,58^, we revisited our annotations of these cells. First, we evaluated their expression of several NK cell related transcripts^50,59^ and found a substantial overlap in gene expression, as demonstrated by a limited panel of genes described previously^60^, that putatively segregates T cells (*CD3D*/*GPR171*/*CXCR6*/*NELL2*) and NK cells (*CX3CR1*/*KLRF1*/*SH2D1B*/*MYBL1*) (Figure 2B). As we described previously^47^, nearly all cells expressed *TRDC*, the delta chain of the T cell receptor (Fig. S2D), and *CD3D* expression was similarly strong, further supporting their annotation as T cells. Nonetheless, it remained possible that there existed some heterogeneity with this cluster, which was covered up by a dominant shared gene expression program.

**Figure 2.**
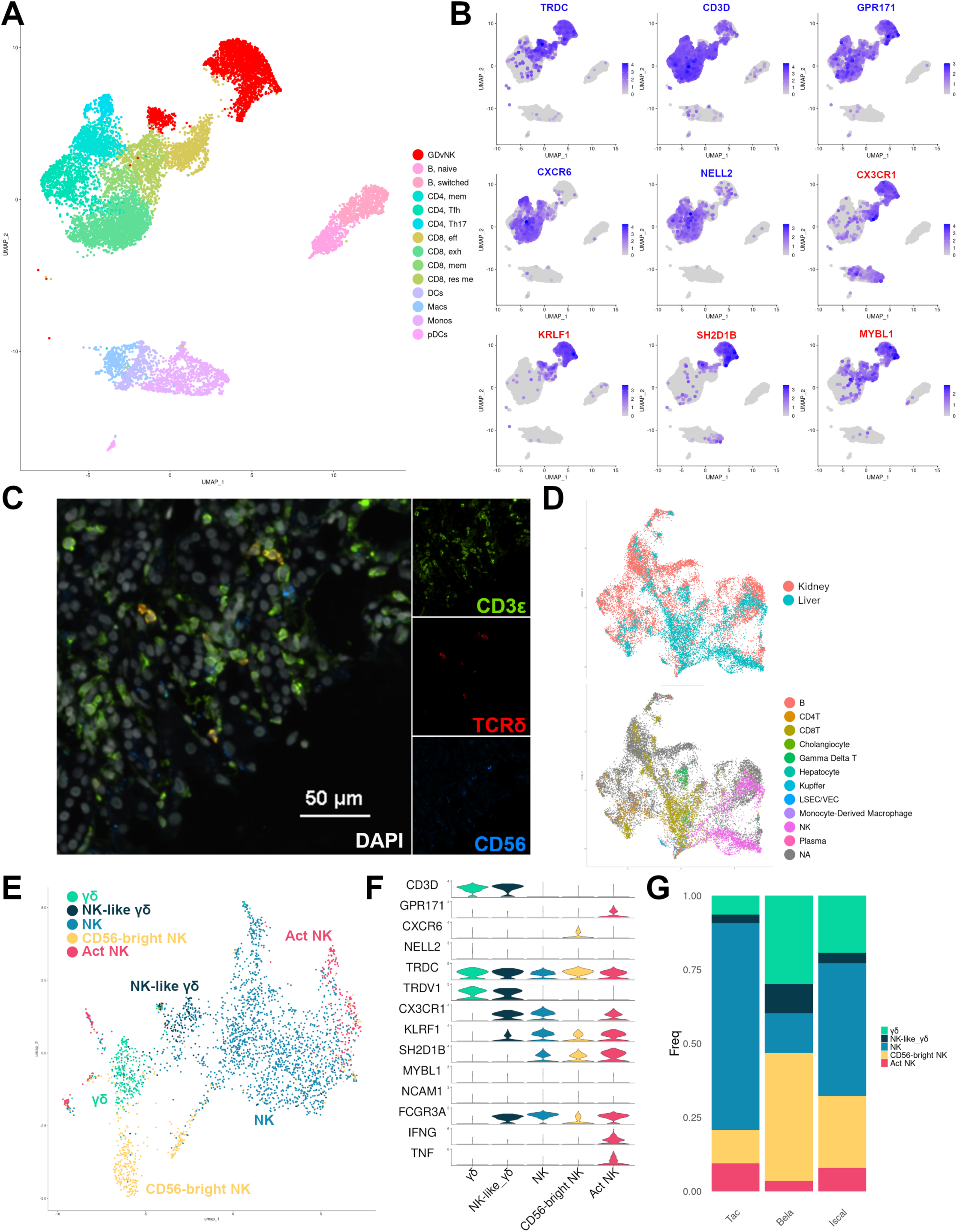
Gamma Delta vs. NK cells in TCMR biopsies. A) UMAP demonstrating distribution of cells previously annotated as γ/δ T cells. B) FeaturePlots demonstrating distribution of indicated genes and lack of segregation of T, NK, and γ/δ T cells, where T cell genes are in blue and NK cell genes are in red. C) Representative immunofluorescent image demonstrating presence of T (CD3ε^+^), NK (CD3ε^-^, CD56^+^), and γδ T cells (CD3ε^+^, TCRδ^+^) cells in TAC_1. D) UMAPs demonstrating successful integration (top) of the Peters and Depasquale dataset, and distribution of those reference markers in the integrated dataset (bottom). E) UMAP demonstrating segregation of annotated γ/δ T cell and NK cell types F) Violin plot showing expression of marker genes G) Bar plot demonstrating distribution of cell type by mIS.

To determine the presence of either or both of these cell types within the allograft, we next performed immunofluorescent staining on four patient-matched clinical FFPE biopsies. Using antibodies against CD3ε, TCRδ, and CD56 we confirmed the presence of T cells (CD3ε^+^ TCRδ^-^), γ/δ T cells (CD3ε^+^, TCRδ^+^), and NK cells (TCRδ^-^, CD3ε^-^, CD56^+^) in all samples (Fig. 2C, S2A). Although limited sample availability precluded large-scale analysis, CD3ε single-positive cells made up the vast majority of identified immune cells in all 4 samples analyzed. Thus, at the protein level, both NK cells and γ/δ T cells are present in similar proportions in rejecting allografts.

Given their presence in FFPE samples, we next interrogated previously published scRNA/CITEseq datasets to further aid in their cellular annotation at the transcriptional level. Analysis of a published CITEseq dataset, which uses barcoded antibodies to combine protein expression with transcriptional landscape, confirmed expression of *TRDC* and *CD3E* in NK cells as well as γ/δ T cells^49^ (Fig S2B). An additional paper used flow cytometric cell sorting to purify a population of CD3ε^-^/CD56^+^ NK cells, and strikingly, our analysis of their data showed that these cells had broad expression of *TRDC*, *CD3E*, and *CD3D* (Fig. S2C)^48^, highlighting the discordance between protein and gene expression and the challenge of using cell surface markers with interrogation of scRNAseq data. Similarly, an published dataset that integrated multiple NK-cell containing datasets demonstrated expression of T cell genes (Fig. S2D) in at least a subpopulation of NK cells, further highlighting the difficulty of separating these populations^50^.

As the prior single cell studies were from peripheral blood, which incompletely capture transcriptional signatures of cells in tissues^48,58^, we next analyzed a dataset, we recently published with our colleagues on TCMR of human liver^61^, wherein NK and γ/δ T cells were more clearly segregated. We successfully integrated the immune cells from our dataset with that dataset (Figure 2D, top), showing clear segregation of NK and γ/δ T cells (Figure 2D, bottom). Using those data as reference, we were able to identify 5 populations of NK and γ/δ T cells in our dataset: two populations of γ/δ T cells, one simply labled “γδ” and one labeled “NK-like γδ”,which expressed NK-cell markers but had clear expression of *CD3D* and *TRDV1*; an “NK” population that likely corresponds to CD16^hi^ cells given expression of *FCGR3A*; a “CD56-bright” population based on lower expression of *FCGR3A* (*NCAM1*, the gene for CD56, is only scantly expressed in these data); and an activated NK population (“Act NK”) that had high expression of both *IFNG* and *TNF* (Figure 2E, F). These cell types were all present across mIS groups, and most patients had every cell type present (Figure S2B). Annotated thusly, NK and γ/δ T cells cluster either separately (NK, Act NK) or near effector and memory CD8^+^ cells (γδ, NK-like γδ, and CD56-bright NK; Figure S2E). Differentially expressed genes between mIS groups was limited (Supplementary Table 2). Use of liver-aided annotation allowed for clearer segregation of these cell types’ marker genes (Figure S2F). These data demonstrate the challenges in annotation of cells from highly inflamed tissues and suggest multi-modal approaches may be necessary to achieve cell annotation.

### 3.3 Class-switched, Activated B cells are Largely Absent in Rejections under CoB

We next focused on intragraft B-cells, in which cellular annotation revealed 6 subclusters of cells, based on differential expression of particular marker genes (Fig. 3A, B)^62^. Three clusters of B cells were identified largely based on their co-expression of *IGHD* and *IGHM*, indicating their relatively immature or naïve state. One very small cluster expressed high levels of *VPREB*, the immunoglobulin ι chain, expressed by pro-B and early pre-B cells. Given their co-expression of *VPREB* with *IGHD* and *IGHM,* we annotated this cluster as preB (Fig. 3A-D). Another small cluster of *IGHD* and *IGHM* expressing B cells also had high expression of *HLA-DQA2*, suggesting recent activation, which we referred to as B_NAIVE, ACTIV_. Both of these small subpopulations were present in patients rejecting under tacrolimus and belatacept but not iscalimab mIS. A much larger population of B cells, present in all patients, also expressed *IGHD* and *IGHM* but lacked markers of activation, and thus are referred to as B_NAIVE_ (Fig. 3A-D).

**Figure 3.**
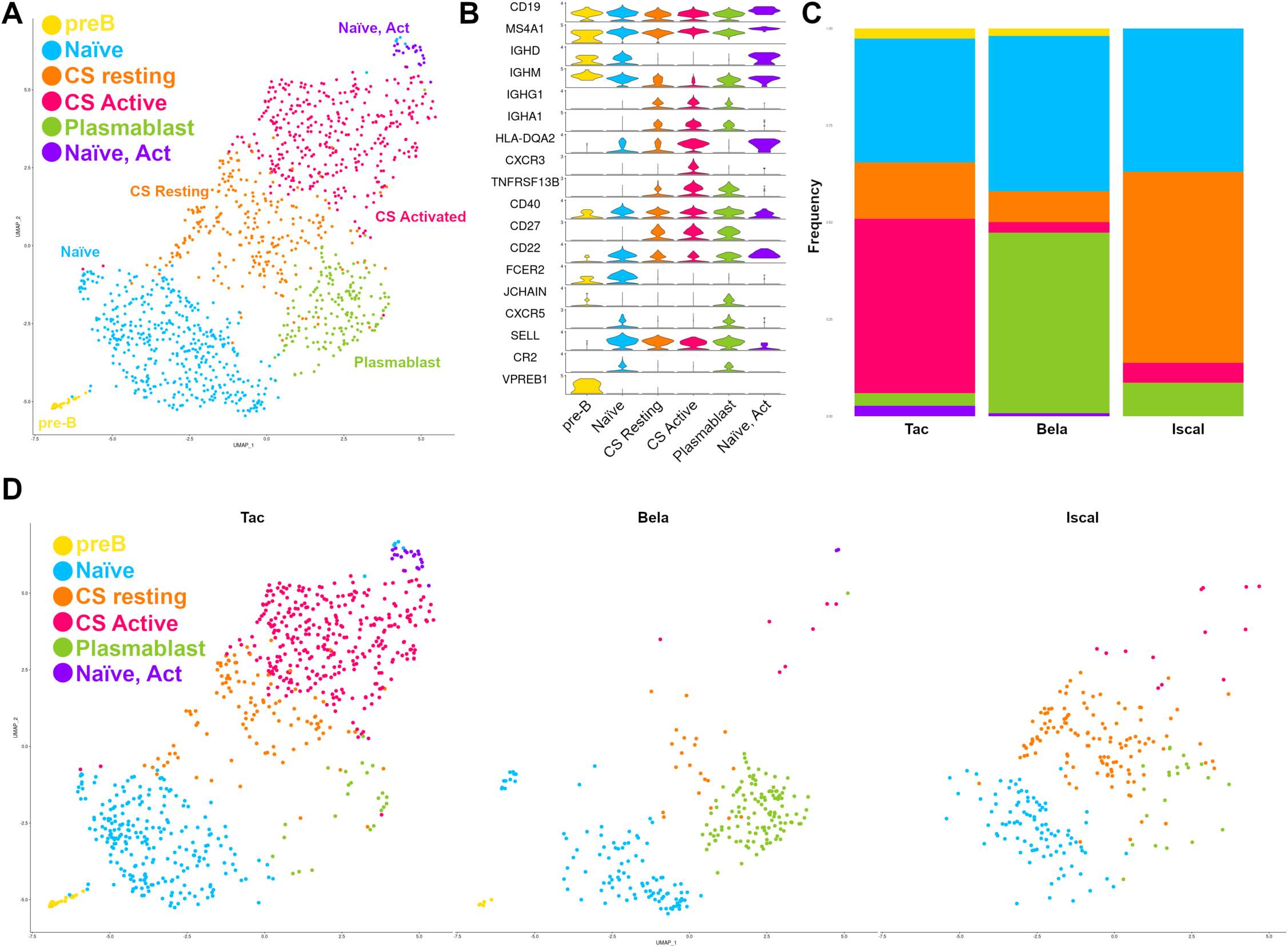
Absence of class-switched activated B cells in grafts from patients treated with costimulation blockade. A) UMAP showing distribution and annotation of identified B cells. B) Violin plot showing expression of subcluster marker genes C) Bar graph showing proportional makeup of B cell populations by mIS regimen. D) UMAP showing distribution of cells separated by mIS regimen. UMAP = uniform manifold approximation and projection; mIS = maintenance immunosuppression; CS = class-switched.

The remaining three clusters all expressed immunoglobulin heavy chain genes indicative of B cell class-switch (CS) and maturation, namely *IGHG1* and/or *IGHA*. We labeled one of these clusters B_CS, ACTIV_ based on its expression of class-switched Ig molecules and the activation marker *HLA-DQA2*. Another CS cluster was labeled as plasmablasts (B_PB_) based on high expression of *IGHM* and *JCHAIN* and lack of expression of *IGHD and PRDM1*^63^. Finally, the remaining cluster was labeled as class-switched, resting B cells (B_CS,REST_) due to expression of *IGHG1* and/or *IGHA* but not *HLA-DQ2* or *CXCR3* (Fig. 3A, B). Intriguingly, only rejections occurring under tacrolimus had a large class-switched populations. The B_CS,ACTIV_ population was rare in all 3 iscalima-treated patients and absent in 2 belatacept-treated patients (Fig. 3C, D, Supplementary Fig. S1C). Furthermore, the B_PB_ cluster was proportionally larger in belatacept treated patients, though this was driven in part by one patient with a very high frequency of these cells (Fig. 3C, D, Supplementary Fig. S2C). Similar to the previous cell types, within-cluster DEG between mIS was limited (Supplementary Table 2). Thus, as expected, CoB substantially inhibited the expansion of class-switched B cells despite their inability to prevent TCMR.

### 3.4 Rejection under CoB demonstrates increased Treg, exhausted populations and decreased peripheral helper CD4^+^ T cells

Given the effect of CoB on emergence of B_CS_ cells, we next analyzed the impact of CoB on the CD4^+^ T cell compartment, initially annotating 8 separate populations (Fig. 4A). First, a clear population of CD4^+^ regulatory T cells (Treg) emerged and was marked by their nearly exclusive expression of *FOXP3* (Fig. 4A, B). Next, we identified a population of exhausted cells based on their high expression of *LAG3, PDCD1, TIGIT,* and *TOX* (Fig. 4A, B). We also identified 2 populations of circulating central memory cells based on their expression of *IL7Ra, S1PR1, TCF7,* and *CCR7* and lack of expression of activation/exhaustion markers (Fig. 4A, B)^64,65^.

**Figure 4.**
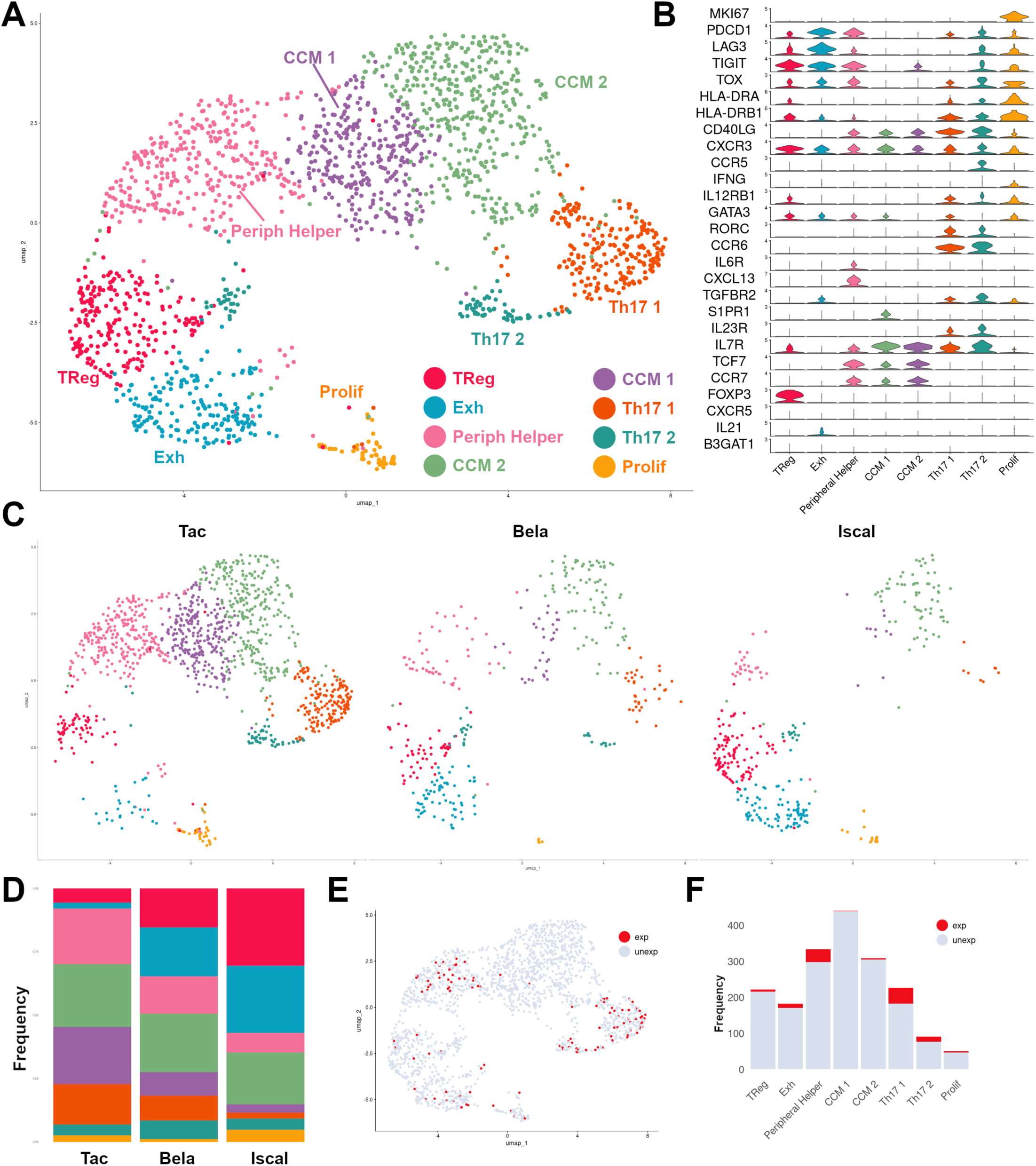
Costimulation blockade is associated with an increase in Treg and exhausted CD4^+^ T cells and decreased peripheral helper CD4^+^ T cells. A) UMAP showing distribution and annotation of identified CD4^+^ T cell populations. B) Violin plot showing expression of subcluster marker genes C) UMAP showing distribution of cells separated by mIS regimen. D) Bar graph showing proportional makeup of CD4^+^ T cell populations by mIS regimen. E) UMAP showing cell types of expanded (exp) and unexpanded (unexp) CD4^+^ T cells. F) Bar graph showing absolute cell counts of expanded (exp) and unexpanded (unexp) CD4^+^ T cells within each subcluster.

Surprisingly, we uncovered two populations of intragraft Th17 cells together defined by their expression of *IL23R*, *RORC*, and *TGFBR2*, but segregated based on differential expression of activation/exhaustion markers *PDCD1, LAG3,* and *TIGIT* (Fig. 4B, Supplementary Fig. S3). Interestingly, we annotated a cluster of CD4^+^ T cells as “peripheral helper” based on their expression of *PDCD1* and *CXCL13* and lower, but detectable levels of *CXCR5* and *BCL6*, consistent with recent work defining a “peripheral” Tfh cells with similar markers (Fig. 4B, Supplementary Fig. S3)^66–69^. Finally, we found a small population of proliferating cells based on their high expression of *MKI67* and an abundance of cell-cycle associated genes (Fig. 4B, Supplementary Fig. S3).

Similar to the results from the B cell compartment, we found different proportions of CD4^+^ T cell types by mIS. Patients treated with CoB demonstrated relatively more cells in the Treg and Exhausted populations than those treated with Tacrolimus, with this pattern being most pronounced in iscalimab patients (Fig. 4C, D, Supplementary Fig. S1D). The remaining cell populations were distributed fairly evenly between mIS types, with noted interpatient variability. (Fig. 4C,D; Supplementary Fig. S1D). Interestingly, within individual clusters, the number of DEGs between mIS was limited, suggesting the effects of mIS are at the population level rather than an impact on gene expression of individual cells (Supplementary Table 2).

We then investigated the clonal expansion of CD4^+^ T cells utilizing scTCRseq identifying individual clonotypes as cells with identical CDR3α/β pairs. Using our prior conservative definition of “clonally expanded” (CD4_EXP_) as clonotypes with greater than 2 cells with identical CDR3αβ sequences^47^, we found relatively few CD4_EXP_ present in the rejecting allografts. Only 7 out of 10 patients had expanded CD4^+^ T cell clones. Within these 7, patients Tac_2 and Tac_4 had 12 and 17 expanded clonotypes, accordingly, for 45 and 55 total cells, respectively, while the others had either 2 (Bela_1) or 1 (Tac_3, Iscal_1, Iscal_2, Iscal_3) expanded clonotypes. Interestingly, the majority of expanded clonotypes were mostly present in the Tph and Th17 clusters (Fig. 4E, F). Thus, although clonally expanded CD4^+^ T cells were far fewer than what we observed for CD8^+^ T cells^47^, they were largely restricted to cells likely involved in the ongoing alloresponse (Tph and Th17).

### 3.5 Costimulation blockade differentially affects expression of costimulatory molecules in select CD4 populations

Although scRNAseq provides substantial transcriptomic information at the level of single cells, it is challenging to resolve cellular interactions from such data. Here, we used computational tools to gain further biologic insights into possible cell-cell interactions, beginning with the CellChat package, which identifies and stratifies cell “senders” and cell “receivers” based on expression of ligands and their accompanying receptors ^54,55^. Given that CoB prevented the accrual of B_CS,ACTIV_ cells, likely based on its ability to inhibit T cell help for B cells, we focused our analysis on potential interactions between CD4^+^ T cells and B cells. As expected, we found differing numbers of interactions in the various mIS backbones (Fig. 5A). In all three groups, Tph cells were important “senders” of cell signals. Interestingly, there was a reduction in total identified interactions under CoB compared with tacrolimus, with a larger effect in the iscalimab group compared to belatacept. An important limitation of this approach is that interactions cannot be assessed for populations that are absent, e.g., interactions between B_CS,ACTIV_ cells and Tph in patients rejecting under Belatacept-based mIS.

**Figure 5.**
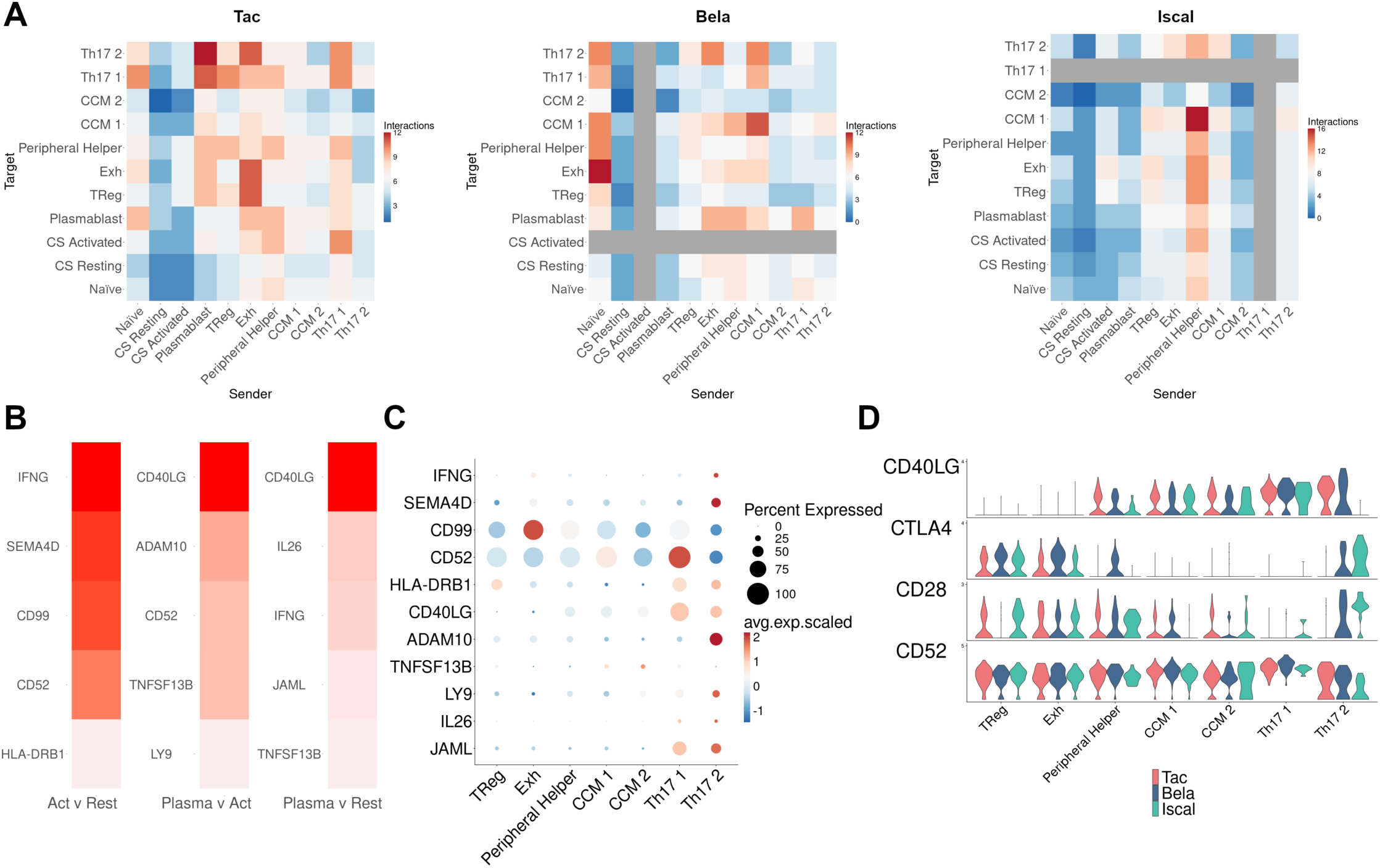
Cellular crosstalk analyses suggest role for peripheral helper and Th17 CD4^+^ T cells and reveal differential expression of costimulatory molecules under different immunosuppressive regimens. A) Heat maps showing the number of non-HLA interactions as determined by the CellChat package between the listed cell types. Each plot shows the interactions identified between the subset of cells treated by the indicated mIS regimen. Interactions unable to be inferred due to lack of a sender or target population represented in grey. B) Top ligands identified by NicheNet package when comparing the indicated recipients (CS activated [Act] vs. CS resting [Rest]; Plasmablast [Plasma] vs CS activated [Act]; Plasmablast [Plasma] vs. CS resting [Rest]) using CD4^+^ T cells as senders. Top 5 ligands as determined by corrected AUPR, with darker color indicating higher corrected AUPR (range 0.012 to 0.038). C) Dot plot showing expression of indicated ligand (y-axis) in the indicated CD4^+^ T cell type. D) Violin plot showing the expression of the indicated ligand in CD4^+^ T cell subclusters, split by mIS regimen. mIS = maintenance immunosuppression; CS = class-switched; AUPR = area under the precision recall curve.

In an attempt to mitigate this limitation, we utilized the NicheNet package, which applies prior models to integrate ligand-receptor and downstream gene expression data to identify interactions that may contribute to gene expression differences given defined sender and receiver populations^56^. By analyzing all three groups of mIS together, possible signaling pathways that contribute to the population differences seen in the mIS groups may be identified. Given the interesting differences in proportions of B cells under different mIS, we used NicheNet to identify ligands that may contribute to changes by comparing three groups: CS Activated vs. CS Resting; Plasmablast vs. CS Activated; and Plasmablast vs. CS Resting; with B cells as the receiver and CD4^+^ T cells as the sender (Fig. 5B). The focus of this analysis was to define genes that may contribute to the different B cell states that might be driven by signals coming from CD4^+^ T cells. Expectedly, *IFNG* and *CD52* were identified as a possible contributors to an activated vs. resting state in B_CS_ cells. Interestingly, consistent with its known role in B cell differentiation, *CD40LG* was identified as the top ligand potentially contributing to differences between the plasmablast cluster and either activated or resting CS B cells. To determine which cells may be contributing those ligands, we then analyzed the expression of the identified ligands by CD4^+^ T cell type (Fig.5C). Notably, we found that the Th17 clusters were the highest expressors of both *IFNG* and *CD40LG*.

As iscalimab directly binds to CD40 and blocks its interaction with CD154^8^, and belatacept binds to B7 molecules and prevents their interactions with CD28^5^, we next examined the expression of these receptor ligand pairs as lower expression of these receptor/ligand pairs has been a mechanism to explain such refractory rejection^19,45,70^. Interestingly, we found decreased of *CD40LG* expression in iscalimab treated patients, specifically in the Tph and Th17_2 populations (Fig. 5D). Under belatacept mIS, *CTLA4* expression was increased in Tph, and CD28 was reduced in Treg cells, consistent with the observation that CD28 can be downregulated upon immunologic stimulation under belatacept^71,72^. Tacrolimus mIS was associated with a decrease in both *CTLA4* and *CD28* in Th17_2 cells relative to the other two therapies. Expression of *CD52* was not notably affected by mIS, nor was ligand expression substantially different on myeloid or B cells (Supplementary Fig. S4).

## 4. Discussion

Acute cellular rejection is a process that is largely driven by donor-specific, CD8^+^ cytotoxic T cells^2^, although many other immune cells contribute to the process. Here we used scRNAseq, which provided unprecedented insights into the transcriptomic phenotypes of diverse immune cell types during TCMR and under the influence of varying mIS. We found the appearance of previously unappreciated γ/δ T cell and CD4^+^ T cell subpopulations, and that the type of mIS contributes to important changes to dynamic shifts in B cell populations. Together, these data describe the trascriptomic landscape of the non-CD8^+^ T cells during TCMR.

One of the more challenging issues we were confronted with was the transcriptional segregation of γ/δ T cells and NK cells. Even in prior studies where investigators purified each cell type using fluorescence activated cell sorting or using CITEseq approaches, clean delineation of these cell types can be difficult, depending upon their environment and cell activation state^48,50,58^. Indeed, our data as well as those from several other datasets, show the substantial overlap of gene expression between NK cells and γ/δ T cells, including *TRDC*. We were initially surprised at this observation, however, prior work has shown that the chromatin in the TCR locus in NK cells is open^48,73^ and that NK cells can undergo rearrangement of the γ and δ chain^74^. Further, our independent analysis of highly purified NK cells showed the presence of *TRDC* transcripts (Figure S2C,D). These data have implications for prior studies identifying “NK” specific transcripts derived from bulk appraoches^60^, as these may not be NK specific, but instead reflect the varying presence of γ/δ T cells and a spectrum of similar cell states spread across more innate (NK), innate and adaptive (γ/δ T), and even adaptive (α/β T) immune cell types. Nonetheless, future studies will be necessary to better distinguish the presence and contribution of both cell types.

The changes within the CD4^+^ T cell and B cell compartment under different mIS regimens is of great interest. First, the presence of Th17 cells within the allograft raises the question of their role and whether they are directly contributory to rejection, as has been suggested^75^, or if they are important only relative to their ability to suppress Treg expansion^41,76^. Indeed, Th17 cells are capable of secreting inflammatory mediators, and gene expression in our study suggests a possible pleiotropic role in supporting rejection through expression of genes such as *IFNG* and *CD40LG*. Interestingly, CTLA4-Ig has been shown to facilitate Th17 development^77^, and expression of *CTLA4* has been shown to confer resistance to belatacept^70^. While we did not see increased Th17 cells under CoB, we did see increased expression of *CTLA4* in Th17 cells, which may offer a potential therapeutic target for rejection under CoB. Perhaps most striking, however, is the relatively large proportion of clonally expanded CD4^+^ T cells within the Th17 compartment, suggesting that, based on our previous work with CD8^+^ T cells in these samples^47^, they are alloreactive and expanding in response to ongoing alloimmune stimuli during TCMR.

Our finding that CoB prevented the accrual of intragraft B_CS_ cells during TCMR is of particular interest. It is well known that CoB can disrupt interactions between Tfh cells and B cells reducing organization of germinal centers inhibiting B cell responses and also that patients under CoB mIS have substantially less DSA and chronic AMR^14–16,78–80^. For example, kidney infiltrating IL-21 expressing Tfh cells promote alloimmunity, DSA development, and treatment with CTLA-Ig decreased numbers of Tfh cells in draining lymph nodes and DSA levels^44^. Our data extend these observations and show that this inhibitory effect of CoB on B_CS_ cells is present during human TCMR. Although this could be operating at the level of TLOs, where intragraft T and B cells have been shown to interact ^81^, we rarely observe TLOs in our TCMR biopsies.

Nonetheless, it is tempting to interpret the presence of Tph and B_CS_ cells in the rejecting allograft as a coupling of cells that will eventually form the underpinnings of future DSA development - a high-risk immunologic scenario. However, this speculation should be tempered by recent elegant work showing that, during chronic AMR, all intragraft B cells analyzed possessed auto-but not specifically not allo-reactivity^40^. One possibility is that B cell populations within allografts are dynamic in terms of their specificity. For instance, early T and B cell infiltrates with significant allo-specificity could promote graft damage which may result in the later accrual of B cells with more specificity to damage associated auto-antigens. Although potentially not specific for allo-antigens, these intragraft B cells may be a surrogate for understanding the developing alloantibody response or lack thereof in patients under CoB. Thus, determining the specificity of intragraft B cells early during TCMR and throughout its resolution would shed significant light on this issue because TCMR is known to predispose to DSA development and chronic rejection^82^. Further understanding the temporal specificity and transcriptomes of intragraft B cells could identify potential therapies to mitigate the development of these graft destructive scenarios. Additionally, the lack of intragraft B_CS_ cells suggests drug efficacy at the level of the B cell, despite its inadequacy at preventing T cell mediated TCMR, supporting a T cell-centric mechanism of resistance to CoB.

Another intriguing finding of our work is the modulation of expression of co-stimulatory molecules based on mIS, a tribute to the plasticity of gene expression within intragraft immune cells. For example, reduced levels of *CD40L* on Tph and Th17 cells in biopsies from rejecting under iscalimab may reflect an inhibition of activation of these cell types given that iscalimab prevents T cell activation^16^. However, this reduction in *CD40L* was not seen in patients rejecting under tacrolimus-based mIS. Although a prior report showed that tacrolimus reduced CD154 on T cells^83^, this was not performed on cells that had “broken-through” mIS. We also saw that *CD28* expression was reduced on Tregs in patients with BRR, which may contribute to allograft rejection as CD28 is important to maintain Treg functionality^84^. Moreover, increased expression of *CLTA4* on Tph in patients with BRR may contribute to the lack of B_CS_ cells. It will be important to disentangle these regulatory networks and how they are impacted by mIS type to avoid potential “whack a mole” scenarios whereby the plasticity of T cell responses leads to escape from mIS and rejection.

This study has several limitations, including a relatively small set of samples and a heterogenous group of patients. Most importantly, however, for future studies, is the lack of spatially-resolved data. During the digestion of biopsy cores for scRNAseq, information regarding the localization of individual cells is lost. We can address this, in part through the use of computational techniques, but those data are inferential at best, heavily reliant upon prior knowledge, and assumptions that gene expression is correlated with protein-protein interactions. The advent of highly-multiplexed spatial technologies including immunofluorescence, mass cytometry, and transcriptomics is beginning to bridge this gap, allowing for detailed analysis of neighboring cells and precise determination of the cell types and cell states present in different neighborhoods throughout the tissue sample^85–87^.

In conclusion, in this study we utilized scRNAseq to analyze the non-CD8^+^ infiltrating immune populations at whole transcriptome scale in kidney allografts at the time of diagnosis of TCMR. In doing so, we identified clonally expanded Th17 and Tph cells and a lack of CS activated B cells under CoB, providing insights into potential mechanisms of resistance and development of subsequent rejections or DSA production. Understanding the unique signatures of all immune cells under various mIS therapies will enable the development of more sophisticated, targeted anti-rejection treatments to facilitate avoidance and resolution of rejection.

## ACKNOWLEDGEMENTS

We graciously thank all the participants, participant families, and other participant support persons that have made these studies possible. We would like to acknowledge the University of Cincinnati (UC) Transplant Research Team and the UC and Christ Hospital Transplant Care Team for their hard work in consenting, following, and caring for research participants. We thank Dr. Meredith Schuh for help with the immunofluorescence protocol. We would also like to thank the members of the UC/CCHMC Center for Transplant Immunology for their guidance and support in this project.

This research was also made possible using the Cincinnati Children’s Single Cell Genomics Core [RRID:SCR_022653], DNA Sequencing and Genotyping Core [RRID:SCR_022630], Bio-Imaging and Analysis Facility [RRID:SCR_022628], and the Bioinformatics Collaborative Serivce [RRID:SCR_022622]. We specifically acknowledge the assistance of Kelly Rangel and Shawn Smith from the Single Cell Genomics Core. This work was supported by a grant from Novartis and from Public Health Service grants AI167482 (T.S.), AI142264 (D.A.H, E.S.W.), UH2AR067688 (D.A.H., E.S.W.), UL1TR000077 (D.A.H., E.S.W.), AI169863 (D.A.H., E.S.W., and B.M.B.), T32DK007695 (J.T.C), KL2TR001426 (J.T.C), and in part by NIH P30 DK078392 Gene Analysis Core of the Digestive Diseases Research Core Center in Cincinnati..

## Abbreviations

AMR: antibody mediated rejection
APC: antigen presenting cell
BRR: belatacept resistant rejection
CNI: calcineurin inhibitor
CoB: costimulation blockade
DC: dendritic cell
DEG: differentially expressed genes
DSA: donor specific antibody
IS: immunosuppression
mIS: maintenance immunosuppression
NK: natural killer
pDC: plasmacytoid dendritic cell
scRNAseq: single-cell RNA sequencing
scTCRseq: single-cell TCR sequencing
TCMR: T-cell-mediated rejection
TCR: T-cell receptor
Tfh: T follicular helper
TLO: tertiary lymphoid organs
Tph: T peripheral helper
Treg: regulatory CD4^+^ T cell
UMAP: uniform manifold approximation and projection

## Supplement

**Figure S1.**
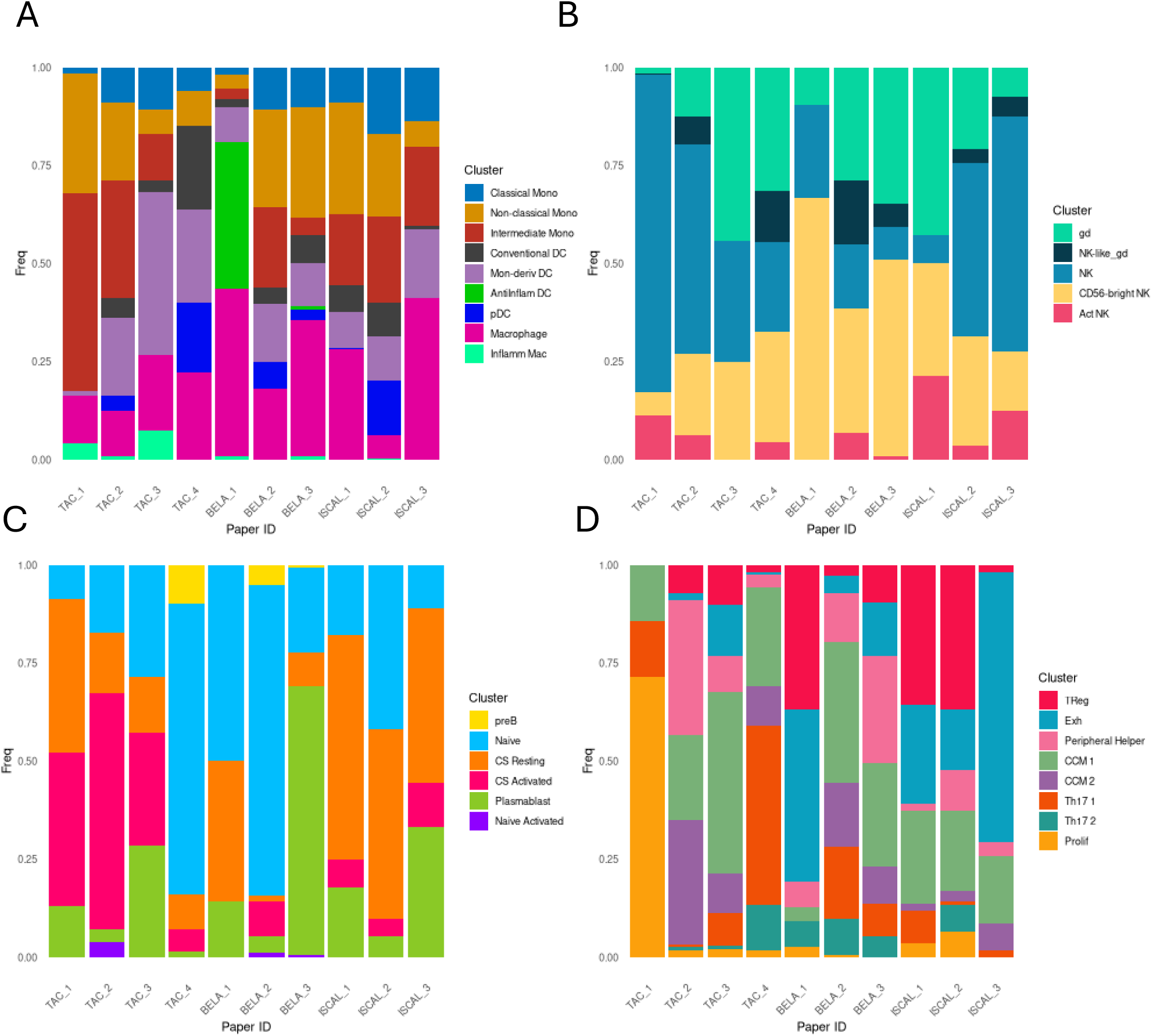
Bar charts demonstrating the proportion of cells in each sample broken down by patient for A) myeloid cells, B) γδ T cells and NK cells, C) B Cells, and D) CD4 T cells

**Figure S2.**
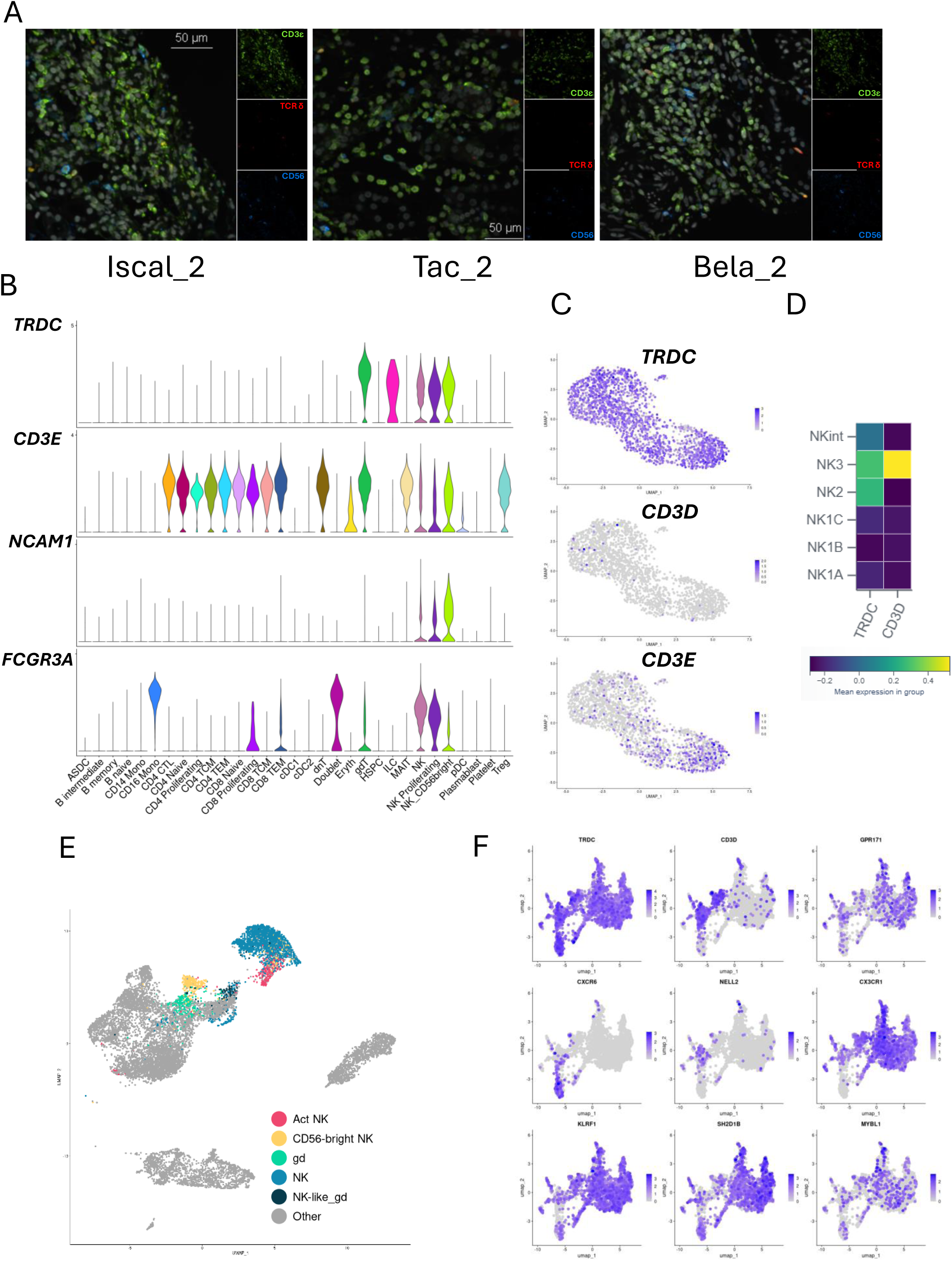
A) Representative immunofluorescence of three remaining patients, demonstrating presence of both intragraft γ/δ T cells and NK cells. B) Violin plots demonstrating overlap of γ/δ T cell and NK cell marker genes in the Hao et al. dataset. C) FeaturePlots demonstrating distribution of γ/δ T cell genes in a sorted NK cell population from Jaeger et al. dataset. D) Plot demonstrating expression of γ/δ T cell genes in NK sub populations from Rebuffet et al. Plot created using the provided web-based data explorer. E) UMAP demonstrating distribution of identified NK and γ/δ T cells on original UMAP space F) FeaturePlots demonstrating distribution and improved segregation of makers on updated NK and γ/δ T cells UMAP space.

**Figure S3.**
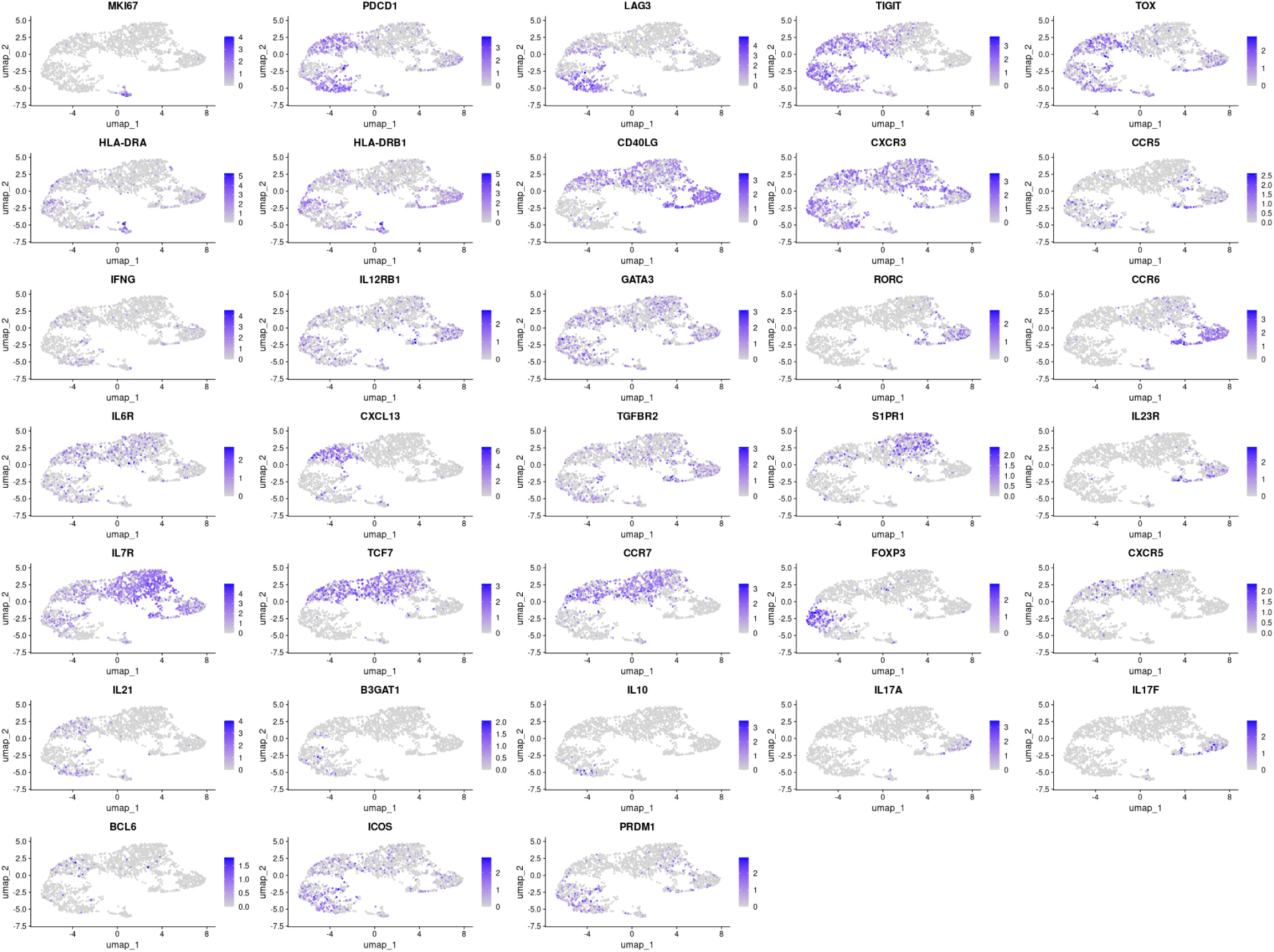
FeaturePlots demonstrating distribution of markers from Figure 3B on CD4^+^ UMAP space

**Figure S4.**
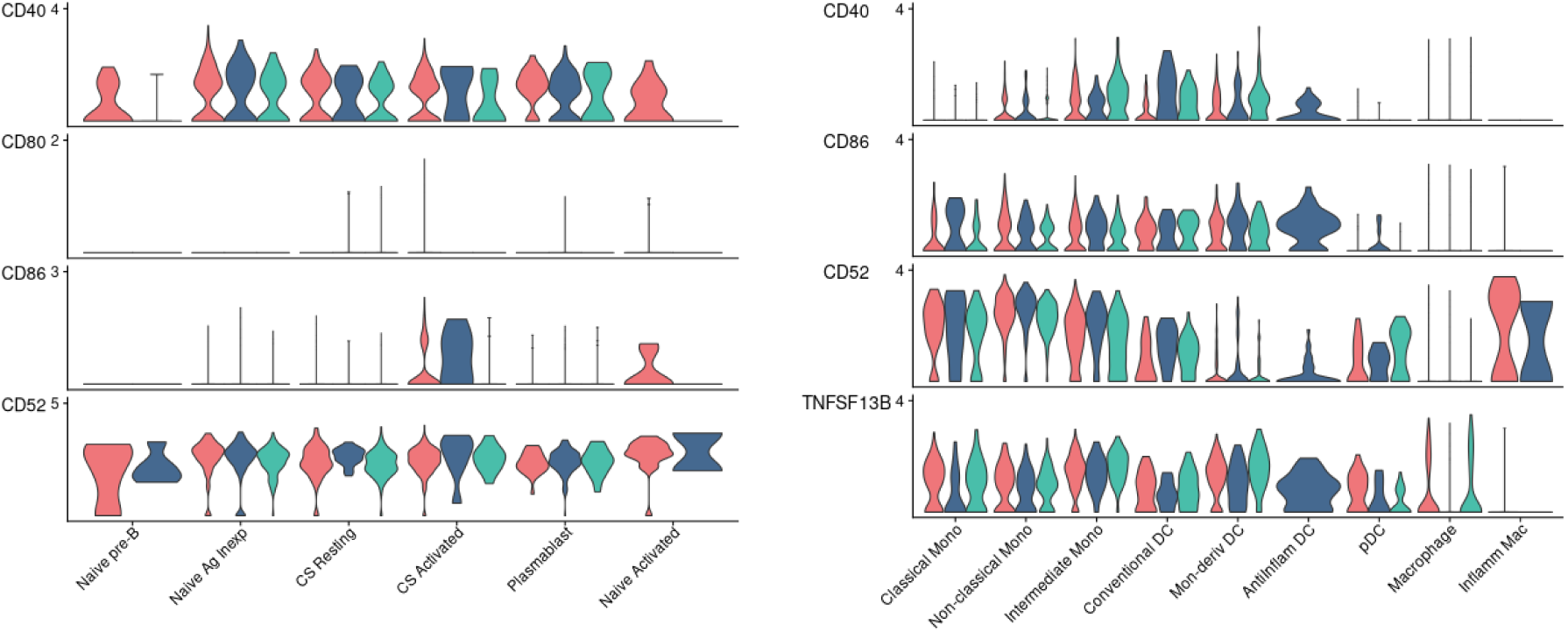
VlnPlots demonstrating expression of indicated ligand genes in B cells (left) and myeloid cells (right)

---

Table S1 DEG for each cluster

Table S2 Within-cluster DEG between mIS for each cluster. Tacrolimus as reference in each case

## Notes

### Competing Interest Statement

The authors have declared no competing interest.

